# Inherited chromosomally integrated HHV-6 demonstrates tissue-specific RNA expression *in vivo* that correlates with increased antibody immune response

**DOI:** 10.1101/741025

**Authors:** Vikas Peddu, Isabelle Dubuc, Annie Gravel, Hong Xie, Meei-Li Huang, Dan Tenenbaum, Keith R. Jerome, Jean-Claude Tardif, Marie-Pierre Dubé, Louis Flamand, Alexander L. Greninger

**Author notes:** These authors contributed equally (Vikas Peddu, Isabelle Dubuc). Co-corresponding authors, Louis Flamand, CHU de Quebec Research Center-Universite Laval, Division of infectious and immune diseases, 2705 Laurier boulevard, Room T1-49, Quebec City, Qc, Canada G1V4G2, Alex Greninger, 1616 Eastlake Avenue East Suite 320, Seattle, WA 98102.

## Abstract

Human herpesvirus-6A and 6B (HHV-6A, HHV-6B) are human viruses capable of chromosomal integration. Approximately 1% of the human population carry one copy of HHV-6A/B integrated into every cell in their body, referred to as inherited chromosomally integrated human herpesvirus 6A/B (iciHHV-6A/B). Whether iciHHV-6A/B is transcriptionally active in vivo and how it shapes the immunological response is still unclear. Here, we screened DNA-Seq and RNA-Seq data for 650 individuals available through the Genotype-Tissue Expression (GTEx) project and identified 2 iciHHV-6A and 4 iciHHV-6B positive candidates. When corresponding tissue-specific gene expression signatures were analyzed, low levels HHV-6A/B gene expression was found across multiple tissues, with the highest levels of gene expression in the brain (specifically for iciHHV-6A), testis, esophagus, and adrenal gland. U90 and U100 were the most highly expressed HHV-6 genes in both iciHHV-6A and iciHHV-6B individuals. To assess whether tissue-specific gene expression from iciHHV-6A/B influences the immune response, a cohort of 15,498 subjects was screened and 85 iciHHV-6A/B^+^ subjects were identified. Plasma samples from iciHHV-6A/B^+^ and age and sex matched controls were analyzed for antibodies to control antigens (CMV, EBV, FLU) or HHV-6A/B antigens. Our results indicate that iciHHV-6A/B^+^ subjects have significantly more antibodies against the U90 gene product (IE1) relative to non-iciHHV-6 individuals. Antibody responses against EBV and FLU antigens or HHV-6A/B gene products either not expressed or expressed at low levels, such as U47, U57 or U72, were identical between controls and iciHHV-6A/B+ subjects. CMV seropositive individuals with iciHHV-6A/B^+^ have more antibodies against CMV pp150, relative to CMV seropositive controls. These results argue that spontaneous gene expression from integrated HHV-6A/B leads to an increase in antigenic burden that translates into a more robust HHV-6A/B specific antibody response.

**Importance:** HHV-6A/B are human herpesviruses that have the unique property of being able to integrate into the subtelomeric regions of human chromosomes. Approximately 1% of the world’s population carries integrated HHV-6A/B genome in every cell of their body. Whether viral genes are transcriptionally active in these individuals is unclear. By taking advantage of a unique tissue-specific gene expression dataset, we show the majority of tissues from iciHHV-6 individuals do not show HHV-6 gene expression. Brain and testes showed the highest tissue-specific expression of HHV-6 genes in two separate datasets. Two HHV-6 genes, U90 (immediate early 1 protein) and U100 (glycoproteins Q1 and Q2), were found to be selectively and consistently expressed across several human tissues. Expression of U90 translates into an increase in antigen-specific antibody response in iciHHV-6A/B^+^ subjects relative to controls. Future studies will be needed to determine the mechanism of gene expression, the effects of these genes on human gene transcription networks and the pathophysiological impact of having increased viral protein expression in tissue in conjunction with increased antigen-specific antibody production.

## Introduction

Human herpesvirus 6 (HHV-6) represents two unique species: HHV-6A and HHV-6B. Primary HHV-6B infection occurs in 90% of children within their first two years of life and causes Roseola, also known as sixth disease, and has been strongly associated with febrile (1). HHV-6B reactivation has been observed in 56% of post hematopoietic stem cell transplant recipients. Those with post-transplant HHV-6B reactivation have also been observed to have a higher chance of human cytomegalovirus reactivation (2).

As with all herpesviruses, HHV-6A/B establish lifelong latency, though it is unique in that it is the only human herpesviruses capable of chromosomal integration. The method of integration is thought to be the result of homologous recombination between the direct repeat (DR) regions on the right end of the HHV-6 genome and the subtelomeric regions of the human genome (3). In approximately 0.5-2% of the general population, integrated virus can be vertically passed through the germline, resulting in one copy of the HHV-6A/B genome in every cell in the resulting child’s body. This is referred to as inherited chromosomally integrated HHV-6A/B (iciHHV-6A/B) (4).

A sensitive ddPCR based assay to detect a 1:1 ratio of HHV-6A/B to human cellular DNA has been described as a method of diagnosing iciHHV-6A/B (5). However, iciHHV-6A/B present a confounding issue for conventional DNA based PCR diagnostic assays when diagnosing HHV-6A/B active infections because HHV-6A/B DNA is always present in the cells of iciHHV-6A/B patients. As a result, though the previously described ddPCR assay can be used to discriminate active infections from HHV-6A/B positive patients, it cannot determine HHV-6A/B active infection in the context of iciHHV-6A/B.

The biological consequences of having the entire HHV-6A/B genome in every cell have yet to examined in detail. In the only large population study performed so far, Gravel et al reported that iciHHV-6^+^ represents a risk factor for development of angina (6). Endo et al also provided convincing evidence of reactivation of iciHHV-6A in a boy with SCID (7). Reactivation was associated with pathogenesis and was successfully treated with antivirals. At the molecular level, the left direct repeat region of the genome is fused to the subtelomeric region (8–10). At the other side of the genome, the right direct repeat ends with telomeric DNA repeat extensions whose lengths are often shorter than those of the other chromosomes (9). In Europe and America, integration in chromosome 17p is overrepresented while overrepresentation of integration in chromosome 22q is observed in Asia (11, 12). Such overrepresentation in chromosome 17p and 22q are unlikely due to multiple independent integration events but likely originates in ancestral integration events that were propagated over time (11, 12).

At present, it is unclear whether integrated HHV-6A/B genomes are transcriptionally active, and if so, whether they exhibit tissue-specific expression. In a recent study Saviola et al reported that the viral genome resides in a condensed nucleosome-associated state with modest enrichment for repressive histone marks H3K9me3/H3K27me3 and does not possess the active histone modifications H3K27ac/H3K4me3 (13). Whether such an epigenetic signature varies depending on the cell type and whether it is modified by *in vitro* passaging of iciHHV-6A/B^+^ subject cells remains to be determined. To study HHV-6A/B gene expression under *in vivo* conditions, we utilized the Genotype-Tissue Expression project (GTEx), which at the time of analysis contained 650 whole blood DNAseq samples, that we used to screen for iciHHV-6A/B individuals. Each of the 650 DNAseq samples corresponds to donor ID-containing RNA-seq data for various tissues within that donor. Here, we report the results of the RNAseq screen, and tissue-based iciHHV-6A/B activity from two unique gene expression datasets. Furthermore, we studied whether the HHV-6A/B gene expression detected was correlated with antigen specific antibody responses. Our hypothesis was that iciHHV-6A/B^+^ subjects may be routinely exposed to a higher antigenic burden than iciHHV-6^−^ subjects and this would translate into a more robust anti-HHV-6A/B immune response.

## Materials and Methods

### GTEx data analysis

GTEx genotype data was downloaded from dbgap (June 1^st^, 2018) using prefetch. 650 DNA sequence SRA files were clipped, decompressed, and extracted using fastq-dump with the following flags: -W (remove tag sequences from dataset), -I (uniquely labels paired-end reads), and -split-files (splits paired-end reads into separate files). The fastq-dump output was piped to bowtie2 (14) for alignment using the --no-unal (unaligned reads not saved in resulting SAM file) and –-local (local alignment) flags. FASTQ files were aligned to the HHV-6A (NC_001664.4), HHV-6B (AF157706.1), and HHV-7 (NC_001716.2) reference genomes with any repeat like regions manually removed.

For the 6/650 files suspected to be positive for iciHHV-6A/B (e.g., >25x average depth across HHV-6A/B reference genomes in DNA-Seq dataset), corresponding RNA-seq FASTQ files for all available tissues (111 total) were downloaded and aligned to HHV-6A/B reference genomes as described above. All reads were confirmed as HHV-6A/B using BLAST with an Evalue<1e-8 against the NCBI nt database (January 5, 2019). For corresponding negative controls, 100 GTEx biospecimen IDs were randomly selected and all available RNA-seq FASTQ files (1903 total) corresponding to those individuals were aligned against HHV-6A/B references as described above.

DNA-seq FASTQ reads corresponding to the six iciHHV-6 positive donors were aligned to portions of the human genes EDAR (NM_022336.4) and Beta-globin (AH001475.2) that were trimmed of human repeats as well as repeat-trimmed HHV-6A (NC_001664.4) and HHV-6B (AF157706.1) reference genomes (Supplement 2), using with the same bowtie2 options specified above. We calculated a normalized depth of coverage by counting the number of reads aligned to the region of interest (R) divided by the total length of the sequence in megabases (B) from the sample and normalizing the highest rpkm from EDAR or beta-globin obtained to 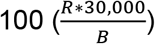.

### GTEx RNA-seq quality control analyses

To calculate fragment insert size distribution of the human WGS and human RNA-seq reads, a random subsample of 10 million reads was taken from GTEx WGS data for each of the four iciHHV-6B positive individuals as well as the 7 RNA-seq samples that showed the highest HHV-6 gene expression. Alignment was performed to hg38 and HHV-6 reference genomes using Bowtie2 with the same flags described above. Because of the low numbers of RNA-seq reads to HHV-6A/B, these reads were pulled from the resulting SAM files and analyzed as one distribution.

HHV-6B RNA-seq reads for each iciHHV-6B individual were aligned to the GTEx-OXRO consensus genome using the Geneious read mapper and manually inspected for variants that could differentiate each donor. Alignment of paired reads to the HHV-6B Z29 reference genome using Geneious was also performed to find paired reads spanning splice junctions.

### Mount Sinai Brain Bank (MSBB) data

Whole exome BAM files aligned to hg19 were downloaded (8/2018) from the Synapse MSBB database (15). Unmapped reads were extracted via samtools using the command “samtools view -b -f 4 <BAM file>” and converted into FASTQ files via samtools bam2fq (16). FASTQ files were then combined and aligned to HHV-6A and HHV-6B reference genomes as described above. Twenty randomly selected samples that were negative for HHV-6 DNA were also screened for HHV-6 RNA. All reads were confirmed as HHV-6 using BLASTn with an Evalue<1e-8 against the NCBI nt database (January 5, 2019).

### Phylogenetic trees

HHV-6A and HHV-6B genomes were downloaded from NCBI GenBank (September 1^st^, 2018). Contiguous HHV-6B sequences between nucleotides 9,515 and 118,889 of the Z29 reference sequence (AF157706.1), corresponding to genes U4 - U77, were used for analysis due to lack of missing sequence in this region. Contiguous HHV-6A sequences between nucleotides 79,352 and 110,248 of the U1102 reference sequence (NC_001664.4) were similarly used, corresponding to genes U48 - U73. Both sets of subsequences were aligned with MAFFT using default parameters. Phylogenetic trees were constructed using the Geneious tree builder with 100 bootstrap iterations.

### Montreal Heart Institute biobank screening

The MHI Biobank contains DNA, buffy coats and aliquots of plasma for every patient (n=15 498). Basic demographic data of the subjects are presented under table 1. For identification of iciHHV-6A/B^+^ subjects, screening was carried out using Roche LightCycler® 480 Instrument II real time PCR apparatus and 384-well plates containing DNA samples from the MHI biobank using a previously described protocol (6). Samples positive for iciHHV-6A/B were confirmed by ddPCR as previously described (17, 18). All patients provided consent and the study was approved by the Montreal Heart Institute Ethics Committee.

**Table 1.**
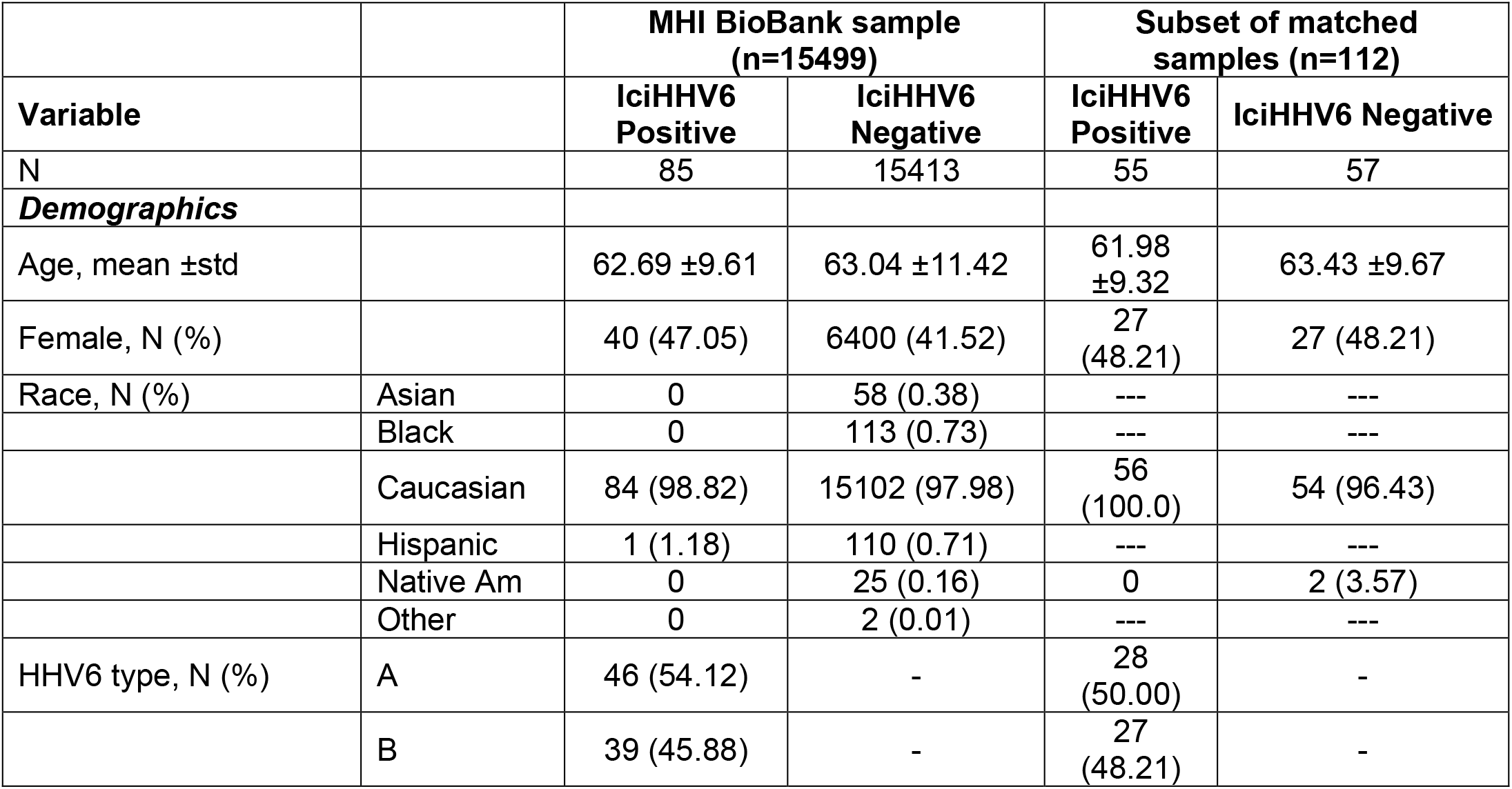
Demographics of the iciHHV-6A/B cohort and control subjects.

### Plasma cytokine analysis

Plasma were analyzed using the Human High Sensitivity T-Cell 14-Plex assay from Eve Technologies (Calgary, Alberta, Canada). Analytes included GM-CSF, IFN gamma, IL-1 beta, IL-2, IL-4, IL-5, IL-6, IL-8, IL-10, IL-12p70, IL-13, IL-17A, IL-23, TNF-alpha.

### LIPS assay

The Luciferase ImmunoPrecipitation assay System has been described previously (19, 20).. In brief, genes of interest are cloned in frame with a FLAG-tagged renilla luciferase (Ruc) gene using the pREN2 vector. All clones are verified by restriction enzyme profiling and DNA sequencing. The following constructs were previously described and generously provided by Peter Burbelo (NIH): pREN2-p18 EBV, pREN2-pp150-d1 CMV and pREN2-HA2 influenza (21). The entire HHV-6B U57 was first cloned into pENTR-FLAG by Gibson assembly cloning. Then, the BamH1/SmaI fragment containing the U57 coding region 1-902 was subcloned into pREN2 to yield the pREN2-HHV-6B MCP. The entire HHV-6B gM (U72 aa 1-344) was cloned by Gibson assembly cloning into BamH1-Sca1 digested pREN2 to yield pREN2-HHV-6B gM. The entire HHV-6B gO (U47 aa 1-738) was cloned by Gibson assembly cloning into BamH1-Sca1 digested pREN2 to yield pREN2-HHV-6B gO. The pREN2-IE1B (aa 1-857) was created by subcloning a BamH1/Kpn1 fragment form the pMalC2-IE1B vector into BamH1/Kpn1-digeted pREN2. The pREN2-IE1A (aa 24- 941) vector was generated by subcloning a BamH1/Xho1 fragment from pcDNA4TO-IE1A into BamH1/Xho1-digested pREN2 vector. The pREN2-GST negative control vector was generated by subcloning the GST gene using the Nco1 blunted-XbaI fragment obtained from pENTR4-GST 6P-1. pENTR4-GST 6P-1 (w487-1) was a gift from Eric Campeau & Paul Kaufman (Addgene plasmid # 17741) (22). The insert was ligated in BamH1 blunted-Xba1 digested pREN2 vector. Protein expression was validated by western blot using anti-FLAG antibodies. To prepare lysates for the LIPS assay, HEK293T cells seeded the day before at 4 ×10^6^ cells/10 cm^2^ petri dish, were transfected with 8 μg of vectors using PEI. Forty-eight hours post-transfection, cells were harvested and lysed as previously described (19, 20). Each serum was tested in duplicate at a final dilution of 1:100. Sera were incubated with the lysate containing 10^6^ relative luciferase unit (RLU) of the target antigen in wells of a 96-well plate for 2h at room temperature with shaking (300 rpm) after which protein A coated magnetic spheres were added to the wells with moderate shaking (300 rpm). After 60 minutes, the plates were loaded onto an automatic ELSA plate washer equipped with a magnetic stand. After 3 washes, the buffer was removed, and the plates loaded into a luminometer with an automatic substrate dispenser (TECAN M200 reader, Morrisville, NC, USA). Light emission was measured over 10 seconds with a 2 sec start delay.

### Statistical analysis

Analyses of cytokine levels were performed using non-parametric Kruskall-Wallis test followed by Dunn’s multiple comparisons test. For antibody levels, statistical significance was determined using the non-parametric Mann-Whitney test. A p value <0.05 was considered significant.

## Results

### Screening for HHV-6 in GTEx DNA-Seq data reveals 6 iciHHV-6 cases among 650 individuals

From the whole genome DNA-Seq data available from 650 GTEx individuals, we determined 6 were consistent with iciHHV-6 – 4 iciHHV-6B and 2 iciHHV-6A. These 6 samples had an average normalized depth of coverage across the HHV-6A/B genome that was approximately half (0.45 ± 0.035) that of human housekeeping genes EDAR and beta-globin, consistent with heterozygous iciHHV-6 at the approximate 1% prevalence typically found in human populations (Figure 1A) (23). Of note, no evidence of chromosomally integrated HHV-7 was found (24).

**Figure 1.**
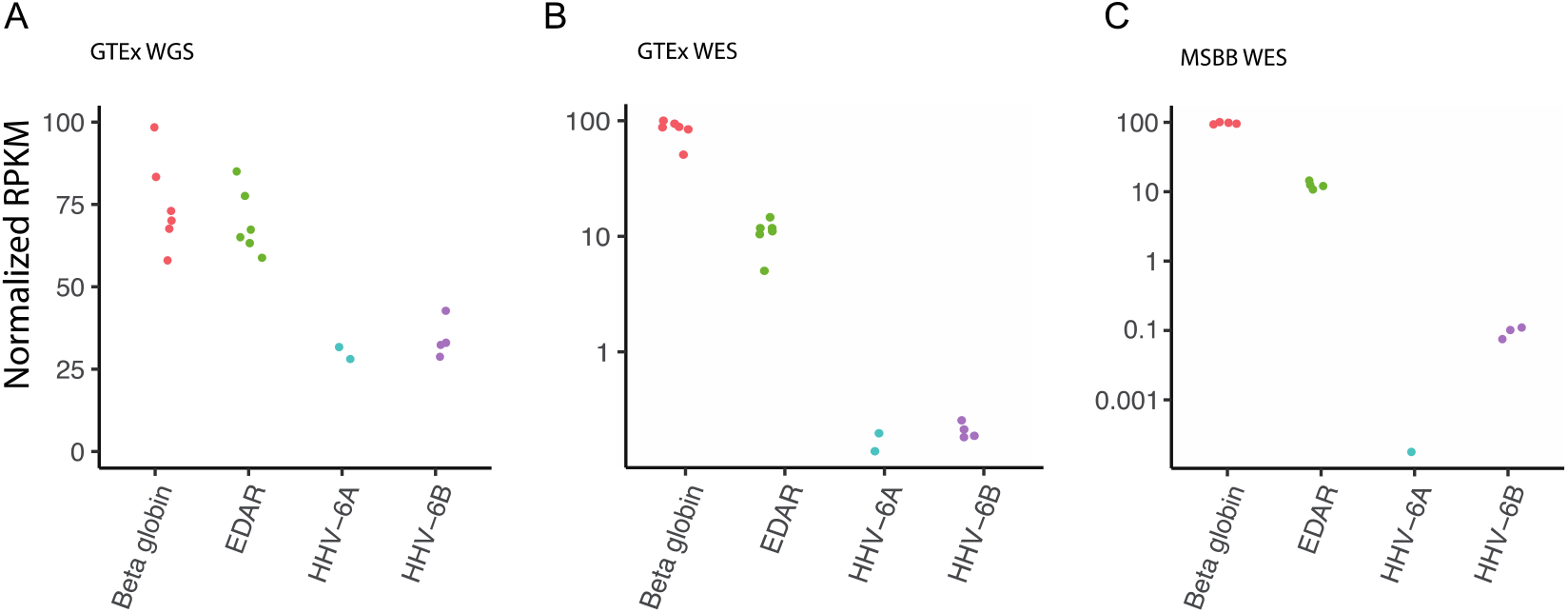
Detection of iciHHV-6A/B individuals in whole genome sequencing (WGS) and whole exome sequencing (WES) data from GTEx and MSBB datasets. a) Six of 650 GTEx samples had high levels of HHV-6A/B in WGS data consistent with iciHHV-6A/B. Normalized depth of HHV-6 compared to control gene reads (EDAR and beta-globin) yielded a ratio of 0.45 ± 0.035, consistent with one copy of iciHHV-6A/B per diploid human genome. b) HHV-6A/B was uniquely detected in off-target reads from the same six individuals’ WES, albeit at far lower levels, consistent with the presence of off-target reads. c) Analysis of the Mount Sinai Brain Bank WES data revealed 4 of 350 individuals were likely positive for iciHHV-6A/B.

### Off-target HHV-6 read coverage from whole exome sequencing also correctly detects iciHHV-6A/B

Because of the unique availability of WGS and whole exome sequencing (WES) data for most participants in the GTEx study, we examined whether WES data could be used to detect iciHHV-6A/B individuals given the comparatively large amounts of WES data available compared to WGS. Within the GTEx WES dataset, the only samples with reads aligning to HHV-6A/B were the six iciHHV-6A/B positive samples by WGS (Figure 1B). The mean depth of coverage for HHV-6A/B in the exome data from iciHHV-6A/B individuals was 0.27x, compared to 117x for beta-globin and 14.5x for EDAR. Screening of the other 603 available whole exome sequences revealed no HHV-6A/B sequence outside of the repeat regions. We used the exome screening approach to screen the MSBB dataset of 350 individuals and found 4 iciHHV-6 positive individuals (1 iciHHV-6A and 3 iciHHV-6B), again consistent with the expected population incidence of iciHHV-6A/B (Figure 1C).

### Phylogeny reveals genetic high genetic similarity amongst iciHHV-6 sequences

Phylogenetic trees reveal clustering with previously deposited iciHHV-6 sequences (Supplementary Figure 1). From the available GenBank HHV-6B sequences, two distinct HHV-6B clades are visible: one Asian and one American/European, with GTEx-OXRO falling into the former, and GTEx-13LV, -14C38, and -YF70 the latter. The American clade consists of iciHHV-6B sequences from the UK and Seattle, as well as HHV-6B sequences from New York (25). The Asian clade contains iciHHV-6B sequences from Pakistan and China, as well as HHV-6B sequences from Japan. It was not possible to build consensus sequences from the Synapse iciHHV-6 exome sequence samples due to insufficient coverage.

### Brain HHV-6 RNA expression is higher in iciHHV-6A individuals compared to iciHHV-6B and glycoprotein U100 and IE1 U90 genes are the most highly expressed HHV-6 genes across tissues

From the six individuals who tested positive for iciHHV-6A/B based on WGS and WES data, RNA-seq data was available from a total of 111 tissues. Analysis of these transcriptomes showed variable tissue-specific activity with highest gene expression levels in U90-U100 genes for both iciHHV-6A and iciHHV-6B (Figure 2; Supplementary Figures 2 and 3). The IE1 protein is among the most divergent protein between HHV-6A and HHV-6B with 62% identity. IE1A and IE1B are large (150 kDa) proteins involved in preventing type I Interferon synthesis and signaling (26, 27). Glycoproteins Q (Q1 and Q2) are part of a multiprotein assembly responsible for receptor binding and viral entry (28, 29). iciHHV-6A expression was noticeably higher in the brain in both the GTEx and MSBB datasets compared to iciHHV-6B (Figures 2 and 3). Viral genes were also actively expressed in the testis and esophagus for both iciHHV-6A and iciHHV-6B individuals.

**Figure 2.**
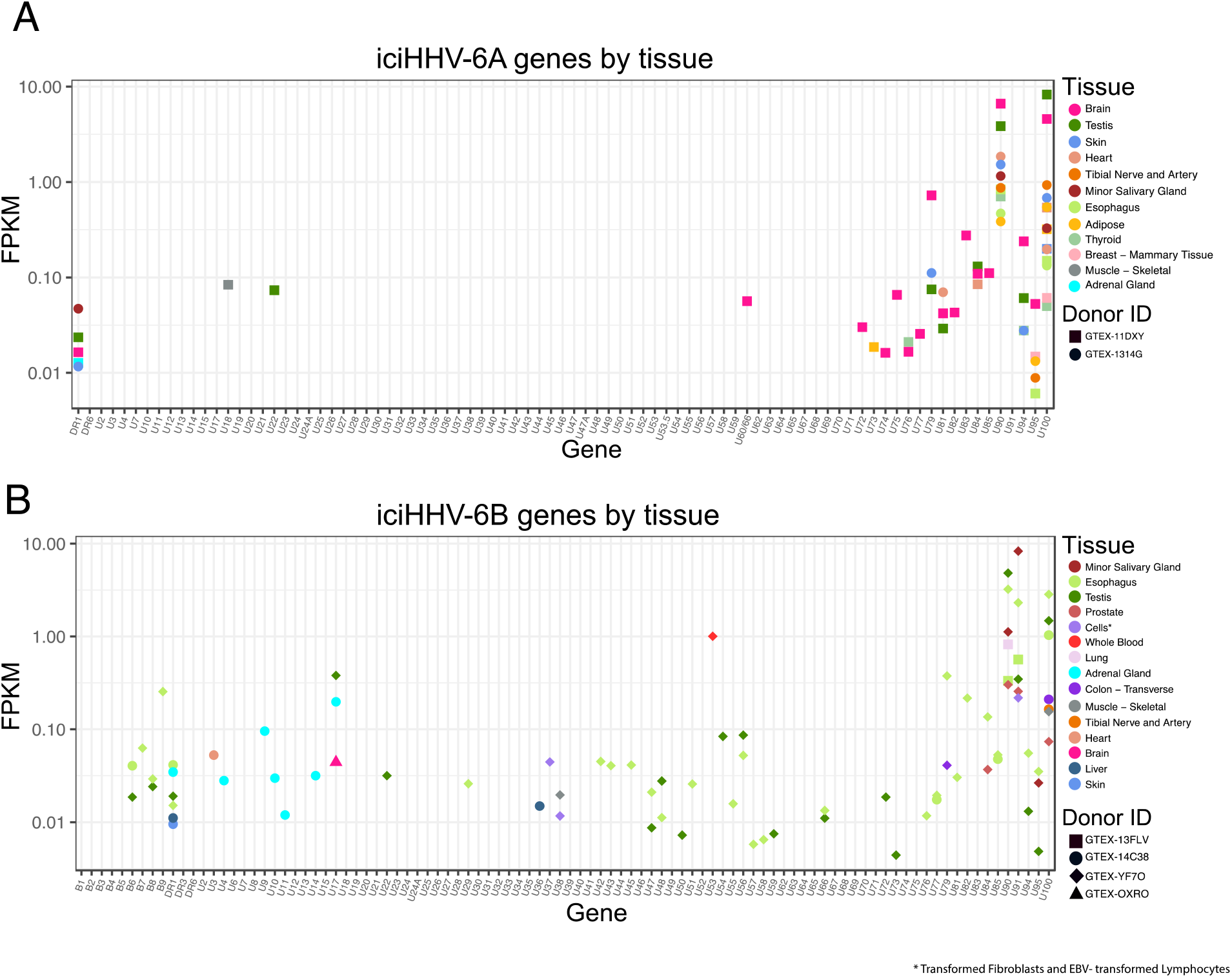
RNA-seq reads from iciHHV-6A (A) and iciHHB-6B (B) positive individuals from the GTEx dataset. The highest levels of HHV-6A/B gene expression were seen in the U90 and U100 genes and in the brain (uniquely for iciHHV-6A), testis, and esophagus.

**Figure 3.**
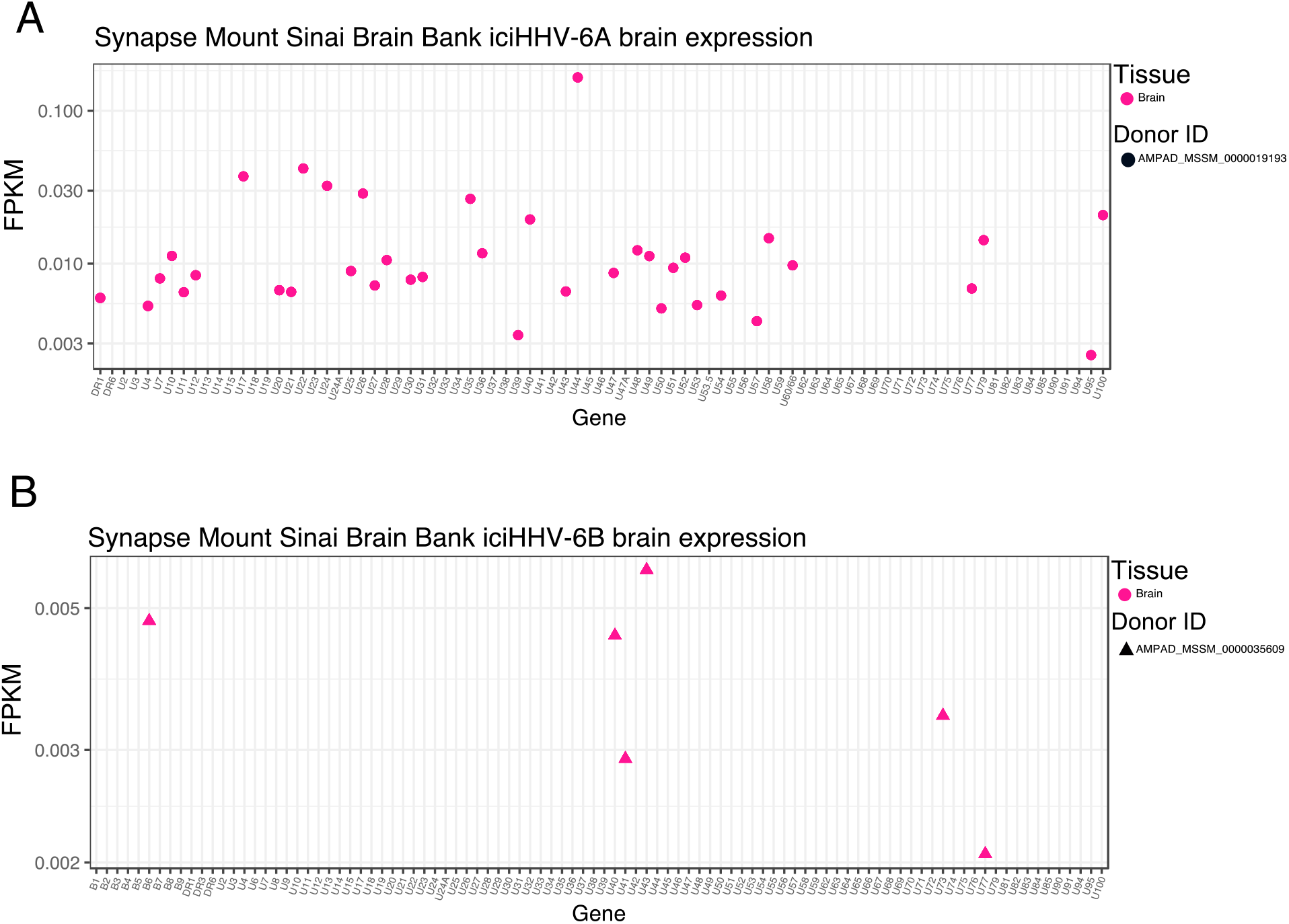
RNA-seq reads from iciHHV-6A (A) and iciHHB-6B (B) positive individuals from the Mount Sinai Brain Bank.

### The presence of reads spanning exons, comparison of SNPs, and insert mean sizes suggest RNA-seq reads are from RNA

Because of the low levels of RNA expression of HHV-6A/B compared to high number of reads to HHV-6A/B in WGS data from iciHHV-6A/B individuals combined with the overall perils of data mining, we performed specific quality control to ensure pre-analytical issues did not compromise our analysis. Specifically, we examined RNA-seq data for fragment insert size, splicing, and presence of unique nucleotide polymorphisms. RNA-seq reads that matched to HHV-6B from iciHHV-6B individuals demonstrated the same fragment insert size as human RNA-seq reads and were noticeably different to the high insert sizes achieved for the human WGS data (Figure 4A). We also detected reads consistent with splicing in the U90 and U100 genes for three of the four iciHHV-6B individuals (Figure 4B). Unique sequence polymorphisms in the HHV-6 RNA-seq data could be found that segregated the four iciHHV-6B sequences and these polymorphisms matched the respective iciHHV-6B DNA sequence for that individual (Figure 4C).

**Figure 4.**
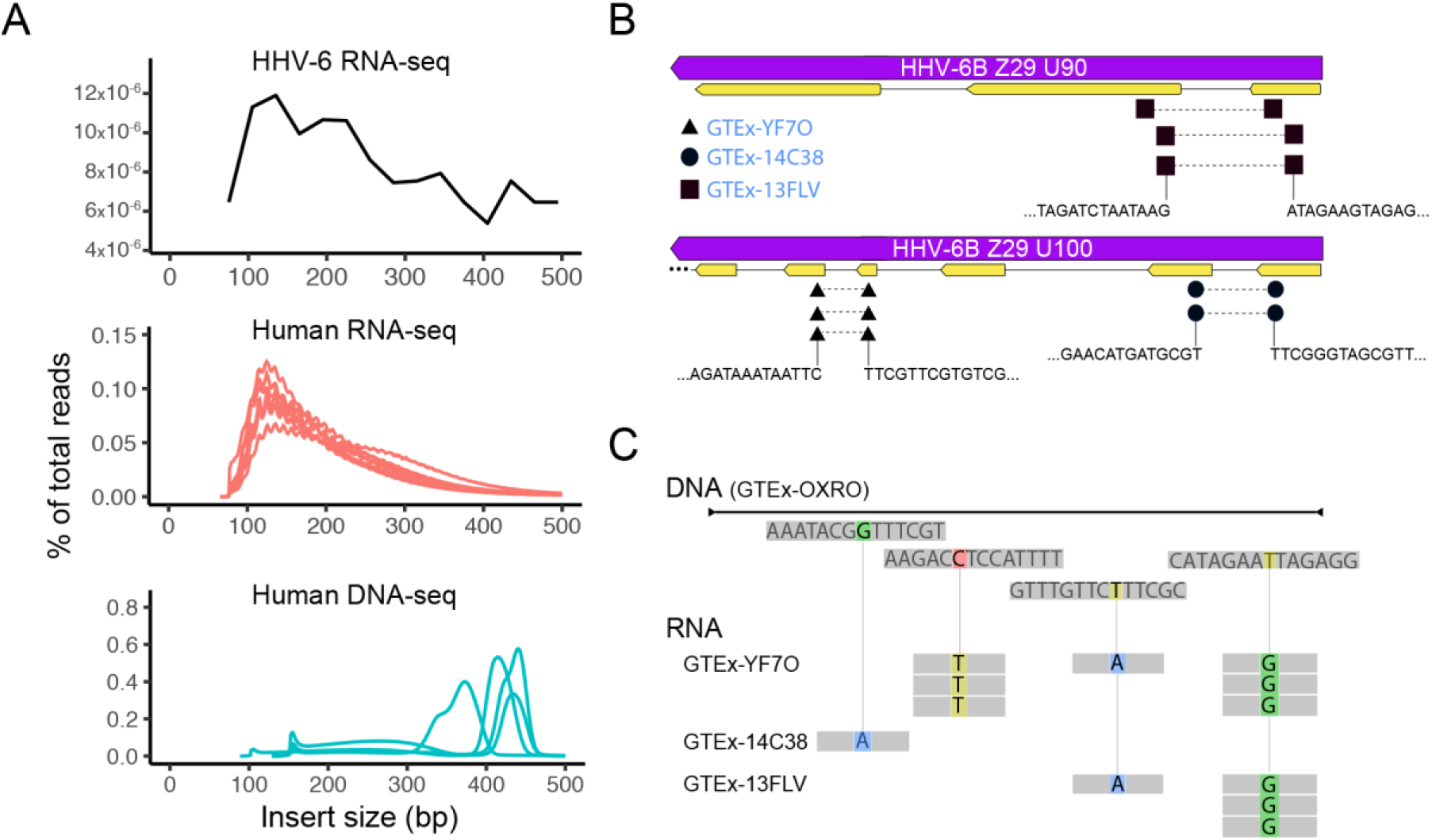
Quality control of RNA-seq reads from iciHHV-6B individuals. A) The insert size distribution for iciHHV-6B RNA-seq, human RNA-seq, and human WGS data were compared to confirm that reads aligning to HHV-6 RNA were not due to HHV-6 DNA contamination rather than RNA. B) Splicing in the U90 and U100 was detected in the three iciHHV-6B individuals with significant RNA-seq reads. C) Sequence polymorphisms in iciHHV-6B RNA-seq reads could also be detected in the three samples that discriminated them from each other and matched their respective consensus iciHHV-6B sequences from WGS data.

### Screening and identification of iciHHV-6^+^ subjects from the MHI biobank

DNA samples from 15498 of the MHI biobank were screen by qPCR to identify iciHHV-6A/B^+^ subjects. In total, 85 iciHHV-6A/B^+^ individuals (40 females) were identified indicating a prevalence of 0.55% (Table 1). Of these, 46 were iciHHV-6A^+^ and 39 were iciHHV-6B^+^. With the exception of two subjects with 2 copies of HHV-6A/cell, all other iciHHV-6A/B^+^ samples were confirmed by ddPCR as having 1 integrated copy of HHV-6A/B/cell. The demographics of the MHI biobank participants are provided under Table 1. Plasma samples from these 85 iciHHV-6A/B^+^ subjects and 20 controls matched for age and sex were used for the serological assay. Blood samples from 55 iciHHV-6A/B^+^ subjects and 57 controls matched for age and sex were used for plasmatic cytokine analyses.

### Plasmatic cytokine analysis

The summary of results is presented under Supplementary Table 1. Of the 14 analytes measured, only TNF-alpha showed a statistically significant difference between controls and iciHHV-6A^+^ subject (p=0.02). All other cytokine levels were comparable between iciHHV-6A^+^, iciHHV-6B^+^ and controls.

### Antibody response of iciHHV-6A/B+ individuals against control antigens

The sera of 46 iciHHV-6A^+^, 38 iciHHV-6B^+^ and 20 controls matched for age and sex were analyzed for their reactivity against FLU (HA), EBV (p18) and CMV (pp150) antigens. The Luciferase-GST fusion antigen was used as negative control. Expression of the fusion proteins was determined by western blot (Figure 5). Relative to the negative control antigen, all sera showed > 2 log_10_ reactivity against the FLU antigen with no significant differences between the groups (Figure 6A). Similar results were observed with the EBV antigen with the exception that iciHHV-6A^+^ and iciHHV-6B^+^ subjects had slightly higher (1.7X) antibody levels relative to control subjects. Such difference did not however reach statistical significance. Results for the CMV pp150 antigen indicate that the cohort contains both CMV seronegative and seropositive subjects (Figure 6C). The proportion of seropositive subjects varied between 25-30% and was no different between groups. Intriguingly, among CMV seropositive subjects, iciHHV-6A^+^ and iciHHV-6B^+^ subjects displayed >5x higher antibody reactivity against the CMV antigen than control subjects with such a difference reaching statistical significance for the iciHHV-6A^+^ group (p<0.01).

**Figure 5.**
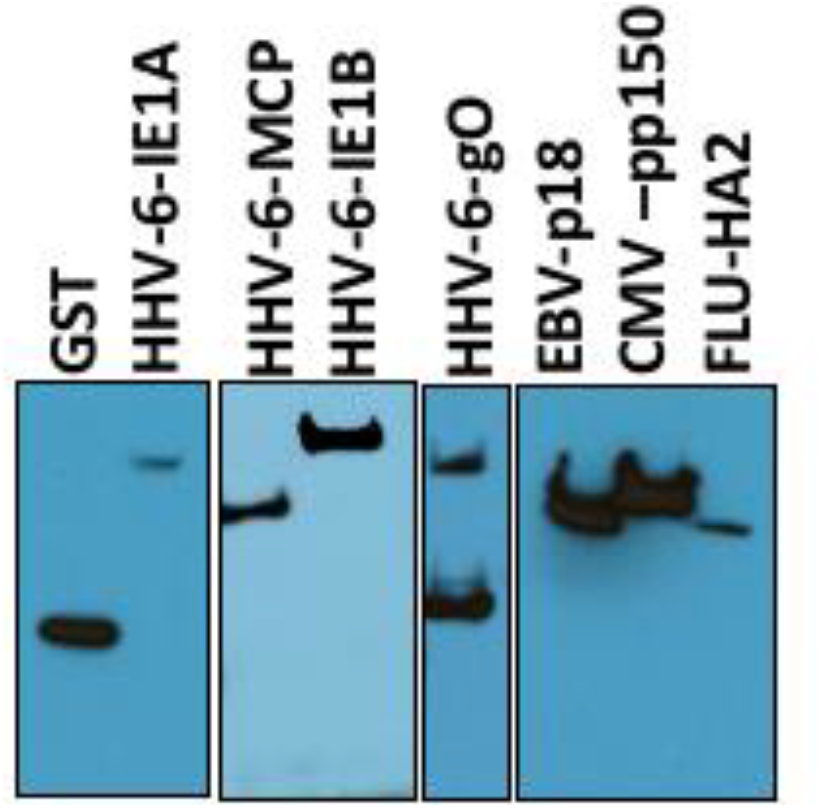
Expression and detection of antigens used for LIPS assay. Vectors expressing control (GST, HA-FLU, p18-EBV, pp150-CMV) or HHV-6 antigens (gO, MCP, IE1A, IE1B) were transfected in HEK293T cells. Forty-eight hours later, expression of proteins was assessed by western blot using anti-FLAG antibodies.

**Figure 6.**
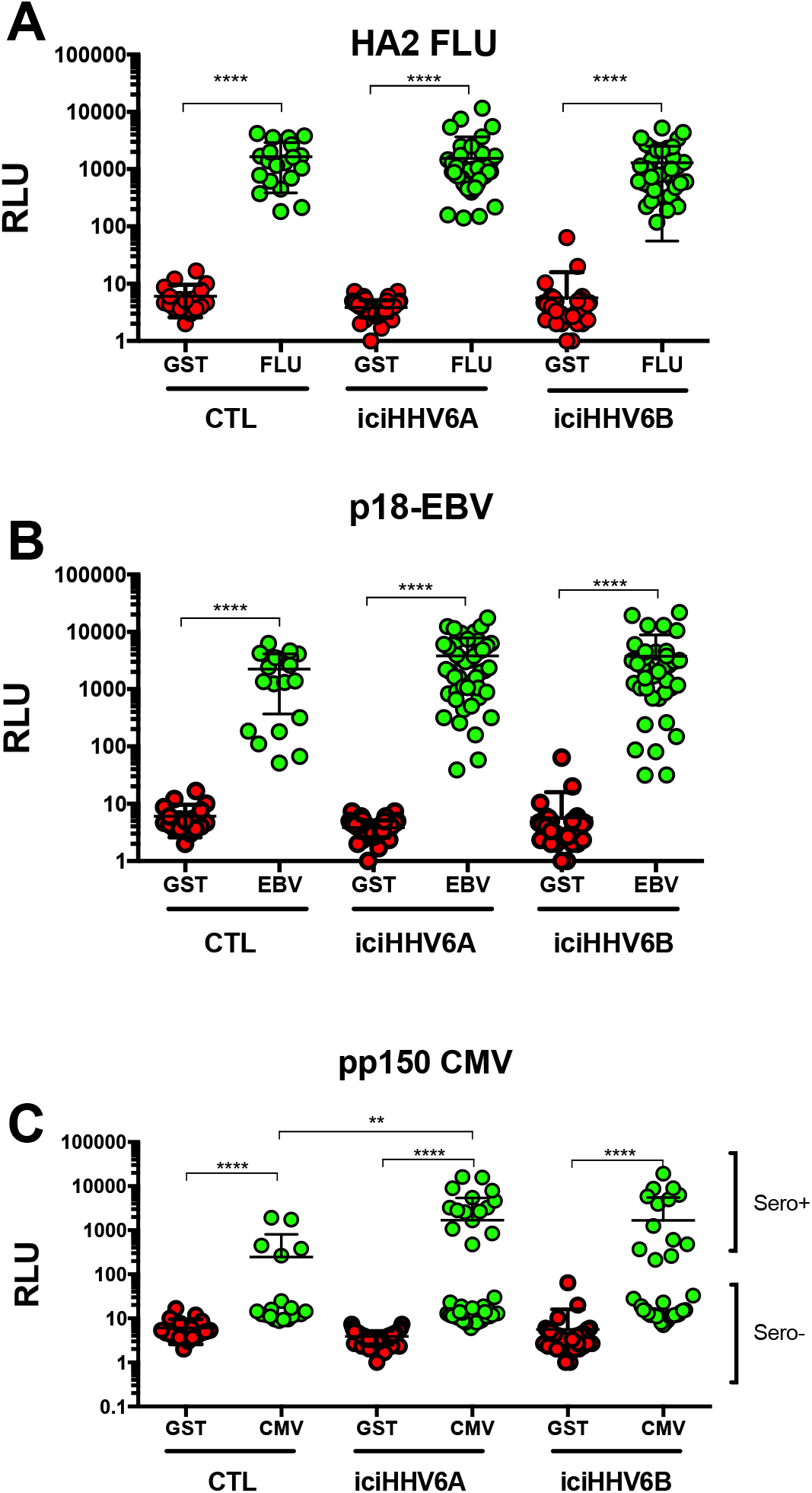
Detection of antibodies against control antigens using LIPS. Lysates containing antigens were incubated with sera from controls (n=20), iciHHV-6A+ (n=46) and iciHHV-6B+ (n=38) subjects to determine reactivity. Each dot represents the mean of samples ran in duplicate from 1 donor. ****p<0.0001, **p<0.01.

### Antibody response of iciHHV-6A/B+ individuals against HHV-6A/B antigens

The GTEx results indicate that certain HHV-6A/B genes such as U90 (IE1) and U100 (gQ) are preferentially expressed relative to others such as U47 (gO), U57 (MCP) or U72 (gM). Whether this would translate into a differential antibody response was examined next. Constructs expressing Ruc-gO, Ruc-MCP, Ruc-gM and Ruc-IE1A or Ruc-IE1B were generated and expression verified by western blots (Figure 5). Most HHV-6A/B proteins share >85% (often >95%) identity at the amino acids level making it very difficult to discriminate whether antibodies are specific to HHV-6A or HHV-6B proteins. For this reason, reactivity against gO (87% identity), gM (97% identity) and MCP (97% identity) were measured using HHV-6B proteins. Sera reactivity against gO, gM and MCP were detected in most subjects with no significant differences observed between iciHHV-6A^+^, iciHHV-6B^+^ and control subjects (Figures 7A-C). Considering that IE1 is the most divergent protein between HHV-6A and HHV-6B (62% identity), antibody reactivity against both proteins were measured. The mean reactivity of sera from iciHHV-6A^+^ and iciHHV-6B^+^ subjects against the IE1A antigen (U90) was significantly higher (p<0.01) that that of sera from control subjects (Figure 7D). Of interest, two distinctive patterns of reactivity were observed in sera of iciHHV-6A^+^ and iciHHV-6B^+^ subjects. Approximately two-thirds of the iciHHV-6^+^ subjects had antibody reactivity against IE1A that was similar to that of the control group. The remaining third iciHHV-6^+^ sera had antibodies levels against IE1A that were ≥25X that of controls. Analysis of antibody response against IE1B also indicate that iciHHV-6B^+^ subjects has significantly more antibodies that control subjects or iciHHV-6A^+^ subjects (Figure 7E). Mean antibody response against IE1B of iciHHV-6B^+^ individuals was 13X greater than that of control subjects (p<0.05).

**Figure 7.**
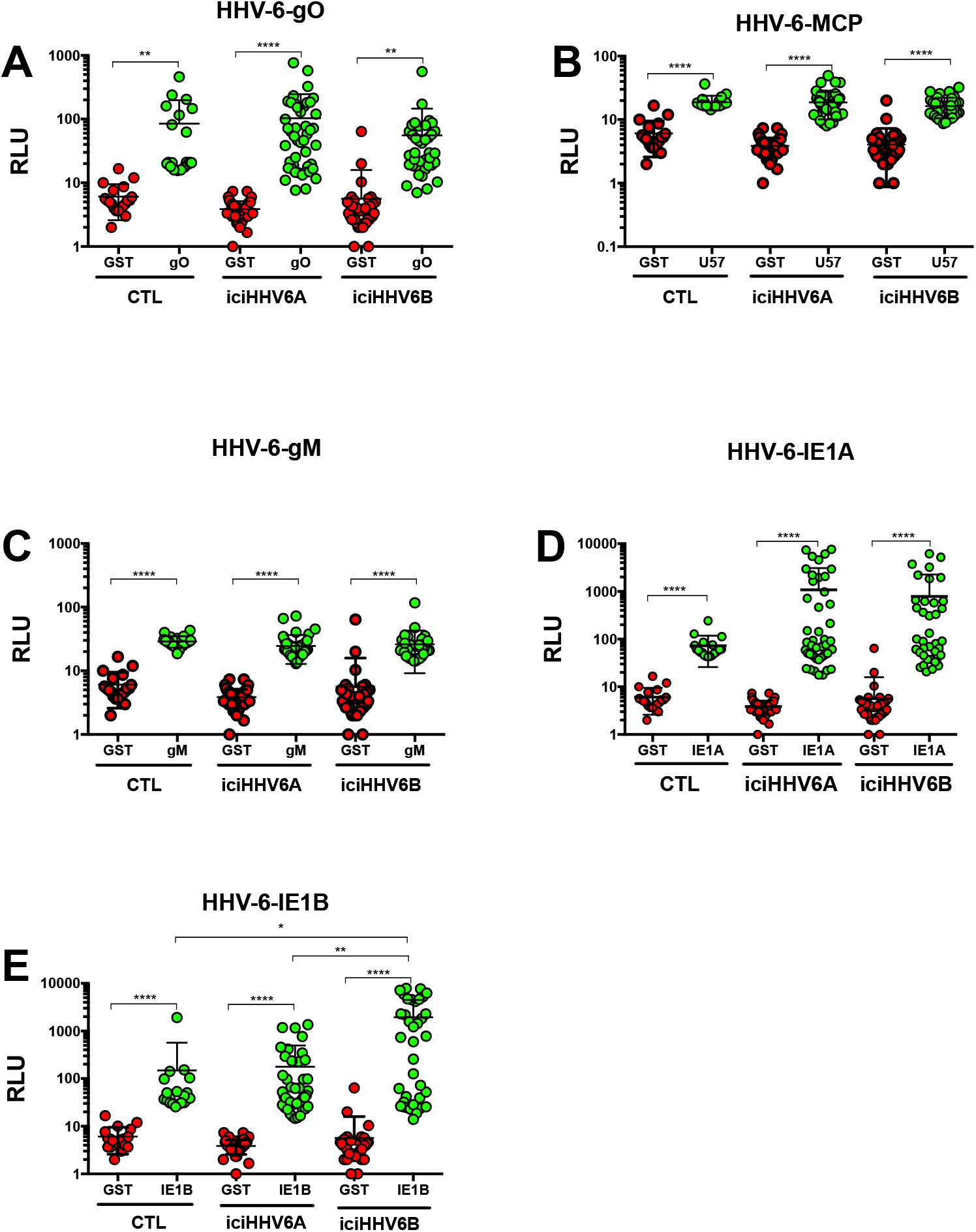
Detection of antibodies against HHV-6A/B antigens using LIPS. Lysates containing antigens were incubated with sera from controls (n=20), iciHHV-6A+ (n=46) and iciHHV-6B+ (n=38) subjects to determine reactivity. Each dot represents the mean of samples ran in duplicate from 1 donor. ****p<0.0001, **p<0.001, *p<0.05.

## Discussion

Here, we show that iciHHV-6 individuals show tissue specific expression of HHV-6 genes in vivo and that the highly expressed U90 gene is associated with a stronger immune response in iciHHV-6A/B^+^ individuals. We detected a 0.92% prevalence of iciHHV-6A/B^+^ in the GTEx cohort, while in the MHI cohort, 0.55% of individuals were iciHHV-6A/B^+^, both matching the previously reported rate of 0.5-2% (30). Our analysis of the GTEx RNA-seq files revealed iciHHV-6A and -6B are not equally active in all tissues, consistent with tissue-specific gene expression. Where HHV-6A/B gene expression could be detected, there was consistently high expression of the U90 and U100 genes relative to other genes for both iciHHV-6A and iciHHV-6B.

We also found that coverage from WES data could prove to be a reliable metric for detecting iciHHV-6A/B in WES sequence. In the GTEx dataset we first analyzed WGS data to find possible iciHHV-6A/B candidates and confirmed the candidates by looking for a 1:2 ratio of coverage for housekeeping genes to HHV-6A/B genomes. Out of the 650 available WES samples in the GTEx dataset, the only ones that contained reads to HHV-6A/B were the six samples confirmed by our WGS screen to be iciHHV-6A/B^+^. Given the significantly greater number of individuals with WES data compared to WGS, the ability to determine iciHHV-6A/B status via WES data alone substantially expands the numbers and types of studies of iciHHV-6A/B status that can be performed.

Within the GTEx iciHHV-6A/B positive samples, brain tissue was observed to be relatively highly active with 11 different viral genes being expressed. We also screened the Synapse Mount Sinai Brain Bank (MSBB) dataset and found three iciHHV-6B sequences and one iciHHV-6A sequence. Analysis of the corresponding RNA-seq reads revealed that iciHHV-6 was expressed in the frontal pole, superior temporal gyrus, parahippocampal gyrus, and interior frontal gyrus. Of the three positive iciHHV-6B samples, only one was found to be expressing HHV-6B genes. This sample, AMPAD_MSSM_0000035609, was only active in the parahippocampal gyrus, and had only 6 genes expressed as opposed to 38 genes expressed for the iciHHV-6A sample (Figure 4). RPKM values for HHV-6B RNA expression in the brain in iciHHV-6B individuals in the MSBB cohort was roughly 2-4 logarithms lower than in other tissues in the GTEx expression data.

Considering that iciHHV-6A/B^+^ individuals have 1 copy of the entire viral genome in every cell, it is expected that at any given time, certain viral genes might be expressed. At present it is unclear what stimulates viral gene expression from the integrated HHV-6A/B. Certain genes such as U90 and U100 are expressed more readily than others, as evidenced by the GTEx data. Of interest, recent results by Gravel et al using *in vitro*-derived cellular clones with chromosomally-integrated HHV-6A also identified U90 and U100 as genes whose expression could readily be demonstrated (18). Expression of U90 and U100 was however clone specific, suggesting that the chromosome carrying the integrated virus might influence viral gene expression.

The biological consequences of carrying iciHHV-6A/B^+^ remain poorly characterized. In the current work, we provide evidence that the magnitude of the antibody responses of iciHHV-6A/B^+^ subjects correlates with the expression of viral genes. The antibody responses against low-level expressed genes such as U47, U57 and U72 are similar between controls and iciHHV-6A/B^+^ subjects. In contrast, the antibody responses of iciHHV-6A/B^+^ subjects against gene products such as U90, which are highly expressed, are much greater than those of control individuals. Whether this increase in antibody against HHV-6A/B antigens offers increased protection against HHV-6 reactivation or reinfection is presently unknown. However, considering that all cells of iciHHV-6A/B^+^ subjects have the potential to express viral proteins, these may, if appropriately presented or exposed to the cell surface, represent targets for immune attacks leading to cell destruction. Over time (decades), such chronic immune destruction may result in various pathological conditions depending on the affected tissues.

Our results also suggest that iciHHV-6A/B^+^ subjects have higher antibody levels than controls against EBV and CMV. Previous work indicated that HHV-6A promotes the reactivation of EBV (31) through activation of the EBV Zebra promoter (32). Immediate-early or early gene products were thought responsible for activation of the EBV Zebra promoter (32). Similarly, a recent report indicated that subjects with documented HHV-6B reactivation had a 15X increase in CMV reactivation rates, suggesting that HHV-6B may, directly or indirectly, trigger CMV reactivation (33). Reactivation of EBV and/or CMV by HHV-6A/B would therefore cause an increase viral antigenic burden resulting in increased antibody production. Considering that every cell of iciHHV-6^+^ subjects contain the entire HHV-6A/B genome, expression of viral transactivator such as U90 may be sufficient to initiate the reactivation of CMV or EBV from latently infected cells.

Since much of the work described here was secondary data analysis, our study has a number of limitations. We were unable to obtain tissues or additional metadata from either the GTEx or MSBB datasets to confirm our work. We cannot rule out the possibility of pre-analytical errors such as trace DNA contamination being the cause of low levels of iciHHV-6 RNA expression. Similarly, we were unable to confirm HHV-6A/B gene or protein expression by orthogonal methods such as immunohistochemistry or RT-qPCR. Alternatively, it is difficult to rule out the possibility that viral gene expression may occur post-mortem, although all GTEx autopsies were performed within 24 hours of death (34). Where possible, we used specific HHV-6 SNPs to ensure no cross-sample contamination could account for recovery of HHV-6A/B reads and we confirmed every read by BLASTn analysis to NT.

Despite these limitations, our work provides the first demonstration of *in vivo* expression of HHV-6 genes from the integrated HHV-6A/B in various tissues. The fact that brain tissue was the most active in transcribing iciHHV-6A genes is of potential interest considering the association of HHV-6A with various neurogenerative diseases such as MS and Alzheimer’s disease (35, 36). Furthermore, these *in silico* analyses could be correlated with actual biological data from a large cohort of iciHHV-6A/B+ subjects. Considering the prevalence of iciHHV-6A/B (0.5-1%), only through analyses of very large biobanks with full medical record will the etiology of diseases linked with iciHHV-6A/B be unraveled.

## Acknowledgements

This work was made possible with grants from the Heart and Stroke Foundation of Canada (LF, JCT, MPD) and the Canadian Institutes of Health Research (LF). We thank the Montreal Heart Institute participants for their contribution to his work.

---

**Supplementary Table 1:** Cytokine levels in plasma of control (n=57), iciHHV-6A^+^ (n=28) and iciHHV-6B^+^ (n=27) individuals.

|  |  | **CTL** | **iciHHV-6A** | **iciHHV-6B** |
| --- | --- | --- | --- | --- |
| **IL-10** | mean | 3,77 | 1,95 | 1,89 |
|  | SD | 11,35 | 1,31 | 1,30 |
|  | p value |  | ns | ns |
| **GM-CSF** | mean | 28,09 | 13,54 | 13,66 |
|  | SD | 81,32 | 12,97 | 11,86 |
|  | p value |  | ns | ns |
| **IFNg** | mean | 11,39 | 9,83 | 8,76 |
|  | SD | 9,53 | 6,33 | 4,83 |
|  | p value |  | ns | ns |
| **IL-12(p70)** | mean | 4,20 | 1,81 | 1,83 |
|  | SD | 16,65 | 1,99 | 1,12 |
|  | p value |  | ns | ns |
| **IL-1beta** | mean | 6,65 | 6,78 | 4,17 |
|  | SD | 10,27 | 8,68 | 6,63 |
|  | p value |  | ns | ns |
| **IL-2** | mean | 7,05 | 23,81 | 3,32 |
|  | SD | 9,31 | 28,71 | 3,04 |
|  | p value |  | ns | ns |
| **IL-5** | mean | 0,74 | 0,57 | 0,56 |
|  | SD | 1,09 | 0,41 | 0,28 |
|  | p value |  | ns | ns |
| **IL-13** | mean | 6,35 | 3,49 | 2,86 |
|  | SD | 20,28 | 4,26 | 2,76 |
|  | p value |  | ns | ns |
| **IL-17A** | mean | 6,60 | 8,16 | 7,85 |
|  | SD | 3,76 | 7,44 | 9,35 |
|  | p value |  | ns | ns |
| **IL-4** | mean | 16,48 | 15,81 | 23,94 |
|  | SD | 22,03 | 27,36 | 48,51 |
|  | p value |  | ns | ns |
| **IL-23** | mean | 397,24 | 422,37 | 322,66 |
|  | SD | 544,71 | 490,85 | 324,10 |
|  | p value |  | ns | ns |
| **IL-6** | mean | 2,28 | 1,82 | 1,11 |
|  | SD | 5,90 | 1,57 | 0,80 |
|  | p value |  | ns | ns |
| **IL-8** | mean | 40,09 | 55,53 | 35,45 |
|  | SD | 68,50 | 60,89 | 60,90 |
|  | p value |  | ns | ns |
| **TNF-alpha** | mean | 8,12 | 11,12 | 6,53 |
|  | SD | 4,00 | 6,00 | 3,72 |
|  | p value |  | 0.02 | ns |

## Supplementary Figures

**Supplementary Figure 1.**
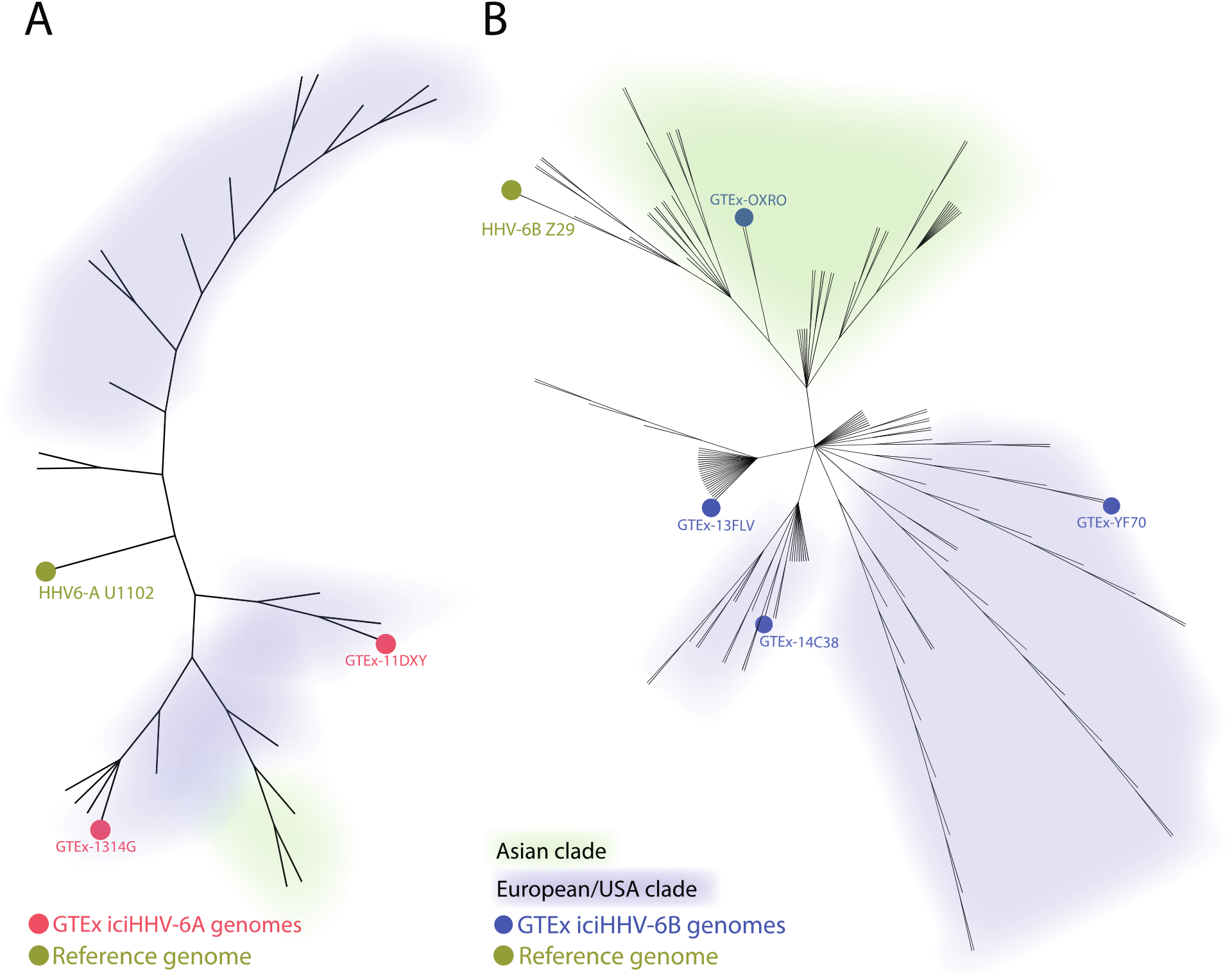
iciHHV-6A (A) and iciHHV-6B (B) sequences found in the GTEx dataset cluster with previously deposited GenBank iciHHV-6 sequences. GTEx-OXRO clusters with a previously deposited iciHHV-6 sequence that is genetically similar to other deposited Asian HHV-6 sequences.

**Supplementary Figure 2.**
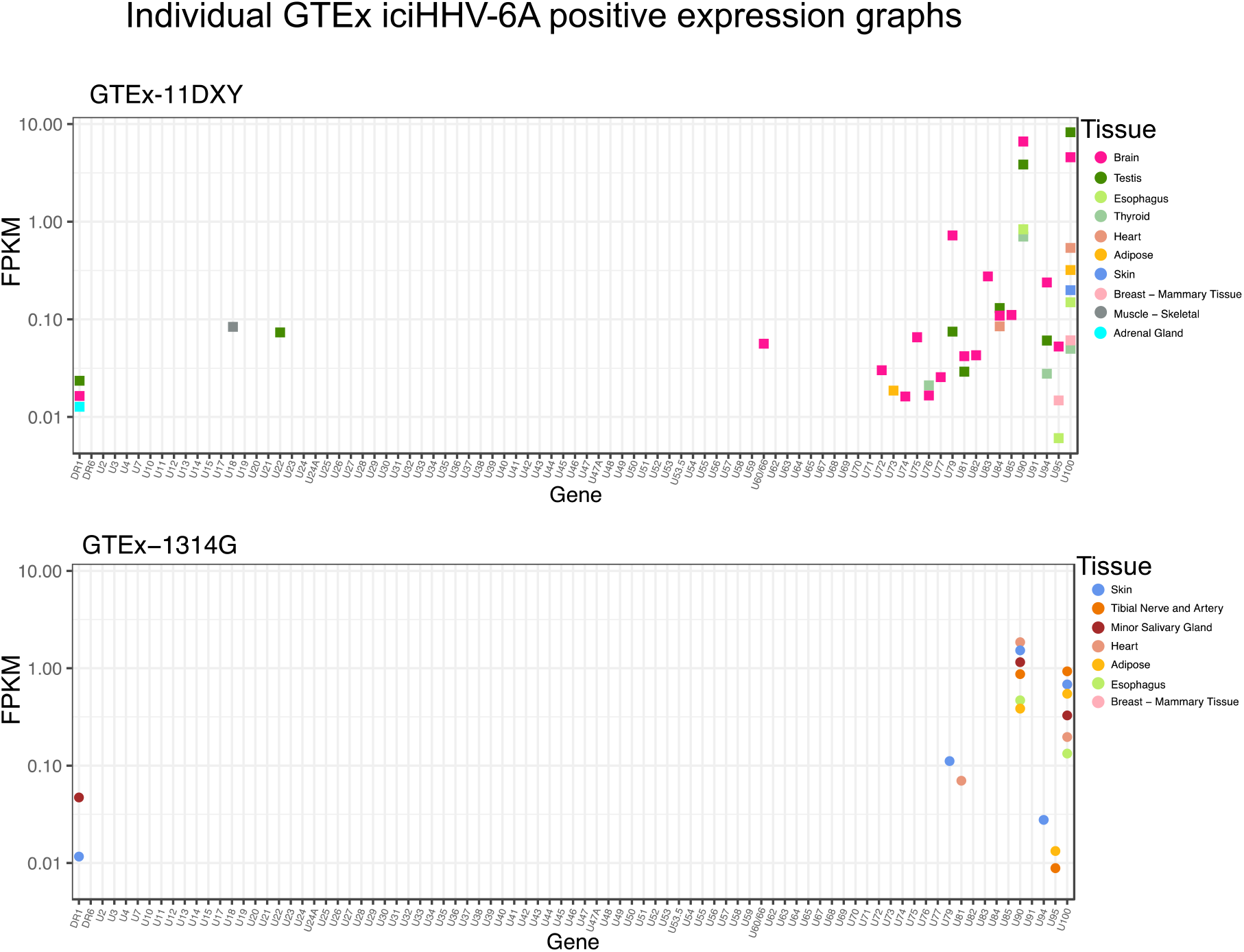
Individual level iciHHV-6A gene expression data from the two iciHHV-6A positive individuals from GTEx.

**Supplementary Figure 3.**
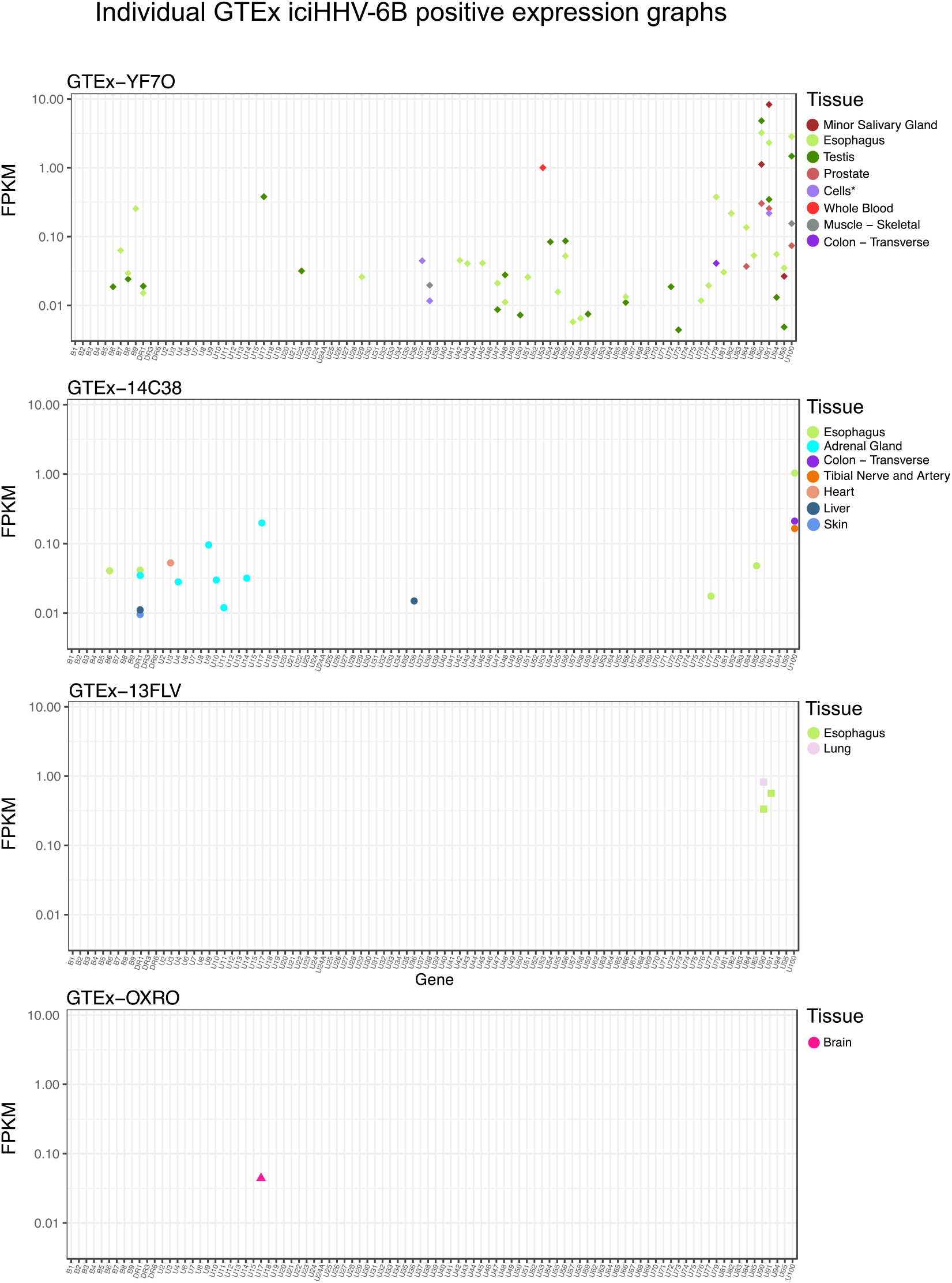
Individual level iciHHV-6B gene expression data from the four iciHHV-6B positive individuals from GTEx.

